# PROSER1 Modulates DNA Demethylation through Dual Mechanisms to Prevent Syndromic Developmental Malformations

**DOI:** 10.1101/2024.07.31.606086

**Authors:** Anna Fleming, Elena V. Knatko, Xiang Li, Ansgar Zoch, Zoe Heckhausen, Stephanie Stransky, Alejandro J. Brenes, Simone Sidoli, Petra Hajkova, Dónal O’Carroll, Kasper D. Rasmussen

**Author notes:** present address: MRC Human Genetics Unit, Institute of Genetics and Cancer, University of Edinburgh, Western General Hospital, Crewe Road South, Edinburgh EH4 2XU, Edinburgh, UK.

## Abstract

The link between DNA methylation and neurodevelopmental disorders is well established. However, how DNA methylation is fine-tuned – ensuring precise gene expression and developmental fidelity – remains poorly understood. PROSER1, a known TET2 interactor, was recently linked to a severe neurodevelopmental disorder. Here, we demonstrate that PROSER1 interacts with all TET enzymes and stabilizes chromatin-bound <u>T</u>ET-<u>O</u>GT-<u>P</u>ROSER1-<u>D</u>BHS (TOPD) complexes, which regulate DNA demethylation and developmental gene expression. Surprisingly, we find that PROSER1 also sequesters TET enzymes, preventing widespread demethylation and transposable element de-repression. Our findings identify PROSER1 as a key factor which both positively and negatively regulates DNA demethylation essential for mammalian neurodevelopment.

## INTRODUCTION

DNA methylation is a fundamental epigenetic process that is essential for normal development. Collaborating with other chromatin-based epigenetic mechanisms, it safeguards the genome by silencing transposable elements, restricts imprinted gene expression, and balances gene dosage between sexes (Jones 2012). Furthermore, DNA methylation can in some cases directly influence transcription factor binding and gene expression through epigenetic modification of promoter and enhancer regions (Schübeler 2015; Rasmussen et al. 2015, 2019; Kreibich et al. 2023). The TET family of DNA demethylases consists of three members (TET1, TET2, and TET3). These closely related proteins contain a conserved catalytic domain that can iteratively oxidize 5-methylcytosine (5mC) to 5-hydroxymethylcytosine (5hmC), 5-formylcytosine, and 5-carboxycytosine, and promote DNA demethylation (Rasmussen and Helin 2016). Mice lacking all three TET enzymes are unable to survive beyond the early stages of development due to gastrulation failures (Dai et al. 2016), while embryos lacking only TET3 can progress to the neonatal stage (Gu et al. 2011). Similarly, although mice lacking either TET1 or TET2 develop normally (Dawlaty et al. 2011)(Moran-Crusio et al. 2011; Quivoron et al. 2011)(Li et al. 2011)(Ko et al. 2011), the combined loss of these enzymes causes developmental abnormalities and increased mortality in a proportion of newborn mice (Dawlaty et al. 2013). The spectrum of developmental defects in knockout mouse lines indicates that TET enzymes have both unique and redundant roles in maintaining developmental processes during early embryonic development.

The function of TET enzymes is modulated via protein-protein interactions with a diverse set of binding partners. For instance, TET enzymes interact with O-linked N-acetylglucosamine (O-GlcNAc) transferase (OGT) (Chen et al. 2013; Vella et al. 2013; Deplus et al. 2013) and the SIN3A-HDAC histone deacetylase complex (Zhang et al. 2015; Williams et al. 2011; Flores et al. 2023; Zhu et al. 2018) to promote histone O-GlcNAcylation and histone deacetylation, respectively. In addition, TET1 and TET2 interact with the two Drosophila behaviour/human splicing (DBHS) proteins Paraspeckle component 1 (PSPC1) and Non-POU Domain Containing Octamer Binding (NONO) (Knott et al. 2016, 2022) to modulate expression of endogenous retroviruses and bivalent genes (Guallar et al. 2018; Huang et al. 2022; Li et al. 2020). Finally, TET2 has recently been reported to interact with Proline and Serine Rich 1 (PROSER1) in the context of UTX and the MLL3/4 branch of COMPASS (Complex of proteins associated with SET1) to modulate H3K4me1 and H3K4me2 levels at UTX binding sites in the human embryonic kidney cell line HEK293 (Wang et al. 2021). The relative importance of these interactions in regulating TET activity and function is an active area of research.

Interestingly, recent findings have linked homozygous loss-of-function mutations in PROSER1 to a novel developmental disorder. This condition features hypotonia, developmental delays, genitourinary malformations, and craniofacial abnormalities associated with sensorineural hearing loss and strabismus (Salah et al. 2022). Given the established role of DNA methylation in neurodevelopment, as evidenced by the association of mutations in DNMT1, DNMT3A, DNMT3B, USP7, and TET3 with a wide spectrum of developmental disorders (Nava and Arboleda 2024), the observed interaction between PROSER1 and TET2 is intriguing. While this interaction suggests a potential mechanism underlying PROSER1-associated syndromes, the precise pathological consequences of PROSER1 deficiency on DNA methylation, gene expression, and ultimately developmental integrity during early embryogenesis remain to be elucidated.

## RESULTS AND DISCUSSION

### Loss of PROSER1 increases preweaning lethality and is associated with developmental disabilities and craniofacial abnormalities

To establish a direct causal link between PROSER1 loss-of-function gene mutations and neurodevelopmental disorders, we generated a mouse line with constitutive inactivation of the endogenous *Proser1* gene using CRISPR-Cas9 gene editing (Fig. S1A, B, C). *Proser1*^+/-^ mice were viable and fertile and these mice were, upon backcrossing to C57BL/6J, intercrossed to generate mice with homozygous PROSER1 loss. Analysis of offspring from these breeders demonstrated that PROSER1 knockout results in partially penetrant pre-weaning lethality (Fig. 1A). We did not observe prominent increases in perinatal lethality suggesting that most PROSER1 knockout embryos may be reabsorbed *in utero* during early gestation. Surviving PROSER1 knockout animals weighed less upon reaching adulthood (Fig. 1B) and displayed frequent eye abnormalities including microphthalmia, anophthalmia, and cataracts as well as intermittent tremors and failure-to-thrive (Fig. 1C, 1D). To further characterise neuroanatomical defects, we performed microcomputed tomography (microCT) scanning of skulls from adult animals (8-14 weeks old). Consistent with reduced overall weight, volumetric analysis revealed reduced cranial bone volume in PROSER1 knockout animals compared to wildtype littermates (Fig. 1E), while bone density remained unchanged (Fig. 1F). Of note, the persistence of this phenotype in fully grown animals suggests that loss of PROSER1 results in permanent developmental disability rather than developmental delays. Further comparison of skull shape revealed distinct malformations of the maxillary and frontal bones as well as a tendency for rounded heads and a general shortening of the skull (Fig. 1G). In summary, germline PROSER1 deficiency in mice results in pleiotropic developmental abnormalities that resemble the human neurodevelopmental disorder in which PROSER1 is mutated.

**Figure 1.**
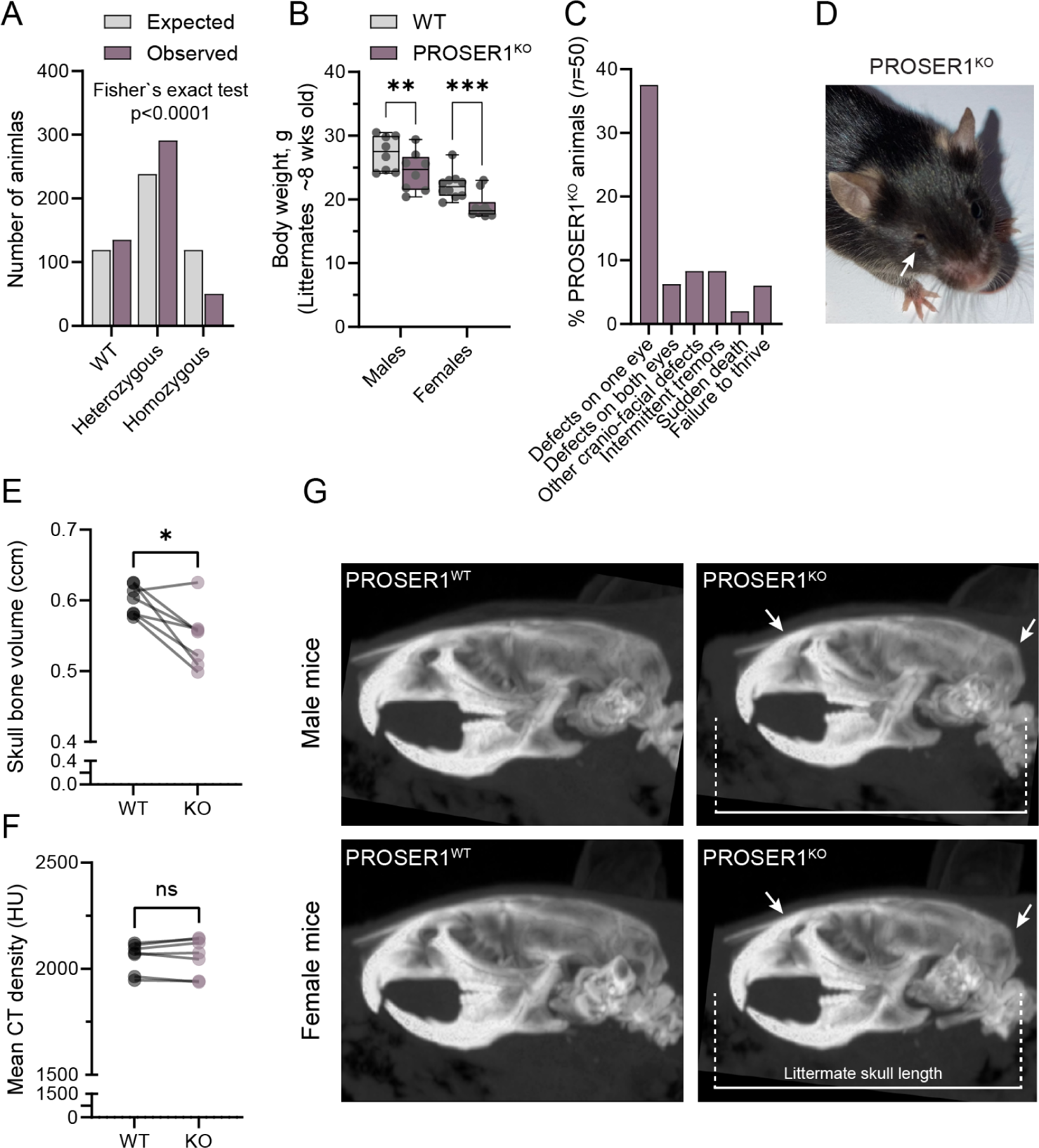
Loss of PROSER1 increases preweaning lethality and is associated with developmental disabilities and craniofacial abnormalities. [A] Histogram showing distribution of genotypes at weaning for 476 pups born from heterozygous *Proser1*^+/-^ breeder pairs. The graph shows expected Mendelian numbers (Expected) compared to the actual distribution of genotypes (Observed). The number of homozygous PROSER1 knockout pups at weaning are demonstrably lower than anticipated, with less than 50% being recovered. Statistical significance for contingency table was measured by Fisher’s exact test (p<0.0001). [B] Box plot showing body weight in PROSER1 knockout (KO) and wildtype (WT) littermates upon reaching adulthood (∼8 weeks of age). Comparisons are made for males and females separately (*n=*8-10). Statistical significance was measured using two-way ANOVA with multiple comparisons (** p<0.01, *** p<0.001). [C] Bar chart showing percentages of gross abnormalities observed in adolescent and adult PROSER1 knockout animals (*n=*50). Eye defects include microphthalmia, anophthalmia, corneal ulcers, and cataracts. Other craniofacial defects include intermittent head tilt and distorted ear (indicative of otitis media) as well as one case of suspected hydrocephalus. [D] Image of a representative PROSER1 KO animal with microphthalmia. The affected eye is indicated by an arrow. [E] Dot and line graph representing volumetric analysis of microcomputed tomography (microCT) scans of PROSER1 KO and wildtype littermate skulls (*n=*7). Statistical significance was measured by paired two-tailed *t* test (* p<0.05) [F] Same as E but representing average CT density (Hounsfield Units) as an indication of bone mineral density. ns indicates not significant. [G] Lateral views of representative microCT scans of a 13-week-old male (above) and 14-week-old female (below) PROSER1 KO animal as well as wildtype littermates. The images are rendered as maximum intensity projections (MIPs) from an equal sized volume to enable direct visual comparison between specimens. Arrows indicate malformations of maxilla and frontal bones as well as rounded head shape. Scale bar indicates skull length of wildtype littermates.

### PROSER1 is a pan-TET interactor and participates in chromatin-associated TOPD complexes

Our data demonstrate functional homology between mouse and human PROSER1, suggesting that mouse embryonic stem cells (mESCs) and their differentiation can serve as an appropriate and tractable model system to elucidate the mechanistic role of PROSER1 in early development. A previous study reported a protein-protein interaction between PROSER1 and TET2 in HEK293 cells (Wang et al. 2021). To determine the conservation of this interaction in mESCs and its potential extension to the entire TET family of enzymes, we raised two specific anti-murine PROSER1 antibodies with epitopes from PROSER1 C- and N-termini, respectively, and performed endogenous IP-MS using PROSER1 knockout mESCs as background control (Fig. 2A, S1D, E, F). Analysis of biological triplicate experiments identified significant enrichment of TET1 and TET2 as well the previously identified TET protein interactors OGT, PSPC1 and NONO. These interactions were also observed in protein lysates isolated from mESC-derived embryoid bodies (EBs). Unlike mESCs, EBs express TET3 and indeed we observed robust enrichment of all three TET enzymes upon PROSER1 IP (Fig. 2B). Importantly, by performing TET2 IP-MS in wildtype and TET2 knockout mESCs we could furthermore recover PROSER1, OGT, PSPC1, and NONO interactions, but no detectable interaction with the other TET enzymes (Fig. 2C). These findings indicate that the presence of TET2 in PROSER1-containing complexes is mutually exclusive with TET1 and TET3, suggesting that PROSER1 forms discrete complexes with each TET protein. Indeed, PROSER1 IP-MS in TET2 knockout cells still robustly enriched TET1 and TET3, demonstrating that their interaction with PROSER1 is not dependent on TET2 (Fig. S2A).

**Figure 2.**
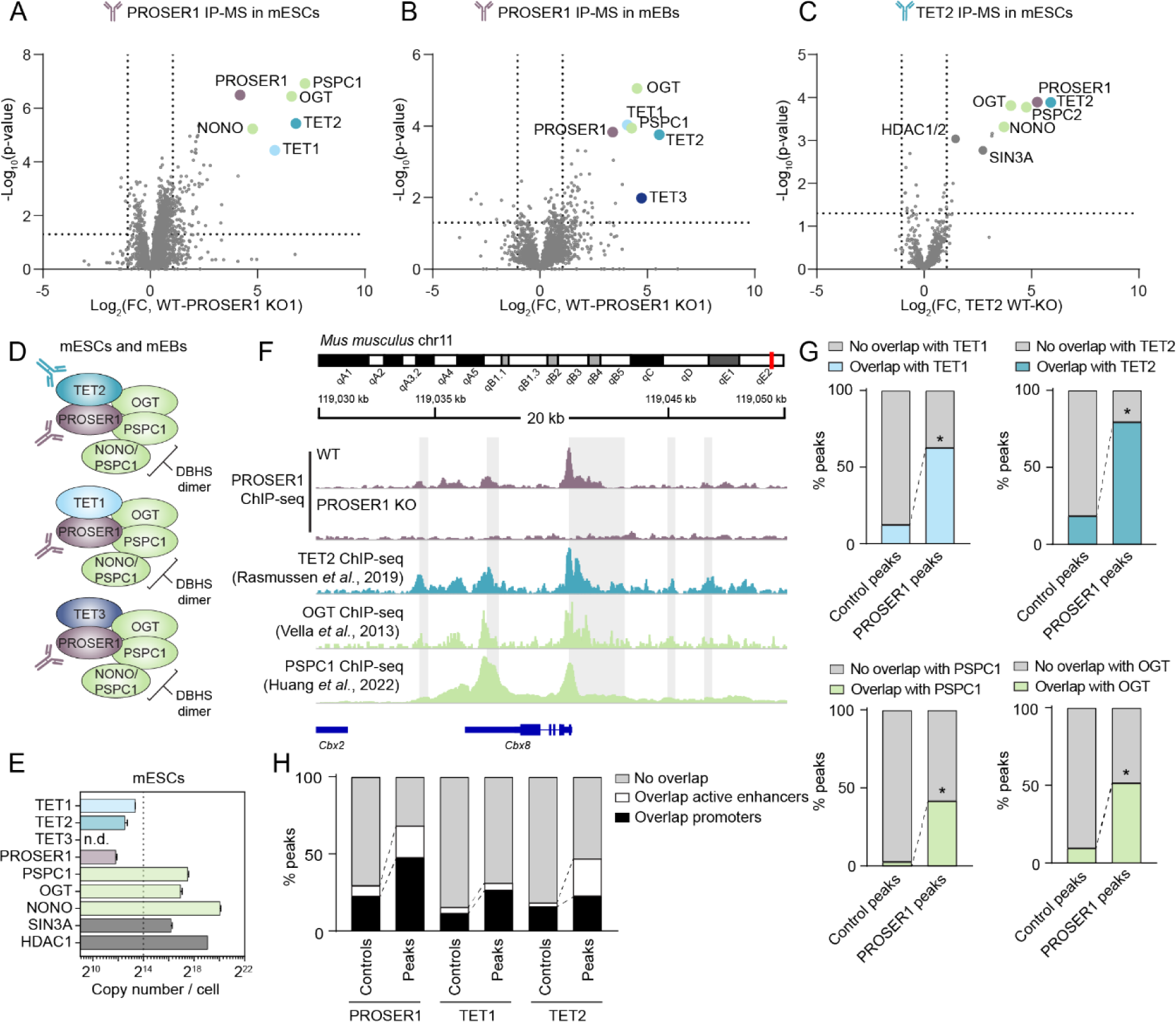
PROSER1 is a pan-TET interactor and participates in chromatin-associated TOPD complexes. [A] Volcano plot showing protein hits from α-PROSER1-C immunoprecipitation and mass spectrometry (IP-MS) in WT mESCs (*n=*3 biological replicates). Protein enrichment is compared to parallel IP-MS in PROSER1 KO mESCs. Dotted lines indicate 2-fold change and p-value 0.05. Proteins of interest are highlighted and identified. See sup. table S3 for full list of enriched proteins. [B] As in A, but carried out in cells differentiated for 2 days to mouse embryoid bodies (mEBs) where TET3 expression is high compared to mESCs. See sup. table S4 for full list of enriched proteins. [C] As in A, but carried out using α-TET2-N antibody and protein enrichment compared to parallel IP-MS in TET2 KO mESCs. See sup. table S6 for full list of enriched proteins. [D] Illustration of the potential composition of TOPD complexes in mESCs and/or mEBs. The DBHS protein dimer is depicted as consisting of PSPC1 and NONO for simplicity but may be variable *in vivo*. [E] Copy number estimation (Wiśniewski et al. 2014) of TET proteins and TET interactors in mESCs determined by MS (*n=*3 biological replicates). The dotted line indicates the sum of TET1, TET2, and TET3 copies, error bars represent the mean ±S.D. [F] ChIP-seq tracks showing a representative region bound by PROSER1, TET2, OGT and PSPC1 in WT mESCs. PROSER1 ChIP-seq in WT and PROSER1 KO mESCs was carried out in this study; sources of the other datasets are indicated in the figure. Protein coding genes in the region are indicated below the tracks. CGIs are indicated in gray. [G] Percentage of PROSER1-N peaks or matched controls (matched to the PROSER1-N peakset in number, size, and distance to DNase hypersensitivity sites in mESCs and generated using Easeq (Lerdrup et al. 2016)) that overlap with TET1, TET2, PSPC1 or OGT binding. All peaksets but PROSER1 were generated from publicly available data (Huang et al. 2022; Rasmussen et al. 2019; Vella et al. 2013; Williams et al. 2011) * p<0.0001, two-tailed Fisher’s exact test. [H] Percentage of PROSER1-N, TET1 or TET2 peaks or matched controls that overlap with active enhancers or promoters.

Our data suggest the existence of multiprotein complexes involving TET proteins, OGT, and PROSER1, as well as members of the DBHS family, which are hereafter referred to as TOPD (<u>T</u>ET-<u>O</u>GT-<u>P</u>ROSER1-<u>D</u>BHS proteins) complexes (Fig. 2D). The relative abundance of these complexes is likely to be affected by variation in the expression of TET proteins and their interactors in different tissues. As mentioned above, TET1 and TET2 are highly expressed in mESCs, whereas TET3 expression is induced in EBs and upon differentiation to neuronal lineages. Similarly, although the DBHS protein PSPC1 robustly associates with TET complexes in all cells tested, PSPC1 can form either PSPC1-PSPC1 homodimers or PSPC1-NONO and PSPC1-SFPQ heterodimers, depending on the relative abundance of each protein (Knott et al. 2016, 2022). Of note, estimation of absolute protein abundance in wildtype mESCs shows that TET proteins and PROSER1 are present at similar copy numbers, whereas the abundance of OGT, PSPC1, and NONO are orders of magnitude higher (Fig. 2E). This implies that excess OGT, PSPC1, and NONO are likely to be involved in processes independent of TET proteins and PROSER1. In contrast, most of the cellular pool of PROSER1 may be engaged within TOPD complexes. Interestingly, PROSER1 IP-MS did not result in enrichment of SIN3A or HDAC1/2 (Fig. 1A, 1B and sup. table S3), suggesting that TET interactions with the SIN3A-HDAC deacetylase complex are independent of PROSER1. We also failed to detect interactions with UTX or members of COMPASS, and profiling of histone modifications by quantitative mass spectrometry revealed little or no global changes in H3K4 methylation in two independent PROSER1 knockout mESC lines (Fig. S1G).

Initial biochemical cell fractionation of mESCs demonstrated that PROSER1 is predominantly a chromatin-associated protein (Fig. S2B). We therefore performed PROSER1 chromatin immunoprecipitation and sequencing (ChIP-seq) using our anti-murine PROSER1-N antibody to gain further insights into its function in chromatin. To ensure specificity of enriched peaks, ChIP-seq was carried out simultaneously on wildtype and PROSER1 KO mESCs. Analysis of biological replicate experiments revealed 1712 high-confidence PROSER1 binding sites (Fig. S2C). Upon intersection with publicly available ChIP-seq datasets in mESCs, we found that a large majority (>95%) are co-occupied by TET1, TET2, OGT, PSPC1 or combinations of these (Fig. 2F and S2D). We next asked if PROSER1 genome co-localisation with each of the complex components is enriched compared to matched control regions. We observed significant enrichment (P<0.0001, Fisher’s exact test) for TET1, TET2, OGT, and PSPC1 co-localisation (Fig. 2G), whereas no enrichment was observed at CTCF binding sites or gene bodies (Fig. S2E, F). Consistent with a role in gene regulation, PROSER1 high-confidence binding sites are associated with both active (H3K27ac and P300) and repressive (H3K27me3 and SUZ12) chromatin domains (Fig. S2G) and overlap regulatory genomic regions such as promoters, CGIs, and active enhancers which are known to be occupied by TET1 and TET2 (Fig. 2H) (Williams et al. 2011; Rasmussen et al. 2019).

### PROSER1 loss disrupts TOPD complexes and alters TET2 genome-wide chromatin binding

To understand how loss of PROSER1 affects the stability of TOPD complexes, we immunoprecipitated TET2 in wildtype and PROSER1 knockout mESC cell lines and analysed eluates by western blotting and label-free mass spectrometry (Fig. 3A, S3A). While PROSER1, OGT and PSPC1, were present in TET2 eluates from wildtype cells, loss of PROSER1 reduced the relative recovery of OGT and abolished TET2-PSPC1 interactions altogether. Furthermore, follow-up analysis of biological triplicate IP eluates using quantitative TMT-labeling demonstrated that TET2-OGT interactions were reduced by ∼60% upon loss of PROSER1, while quantitative recovery of SIN3A was not affected (Fig. 3B). Of note, we found that the level of TET2 O-GlcNAcylation was unchanged in PROSER1 knockout cells, suggesting that enzyme-substrate interactions between TET2 and OGT are preserved in the absence of PROSER1 (Fig. 3A).

**Figure 3.**
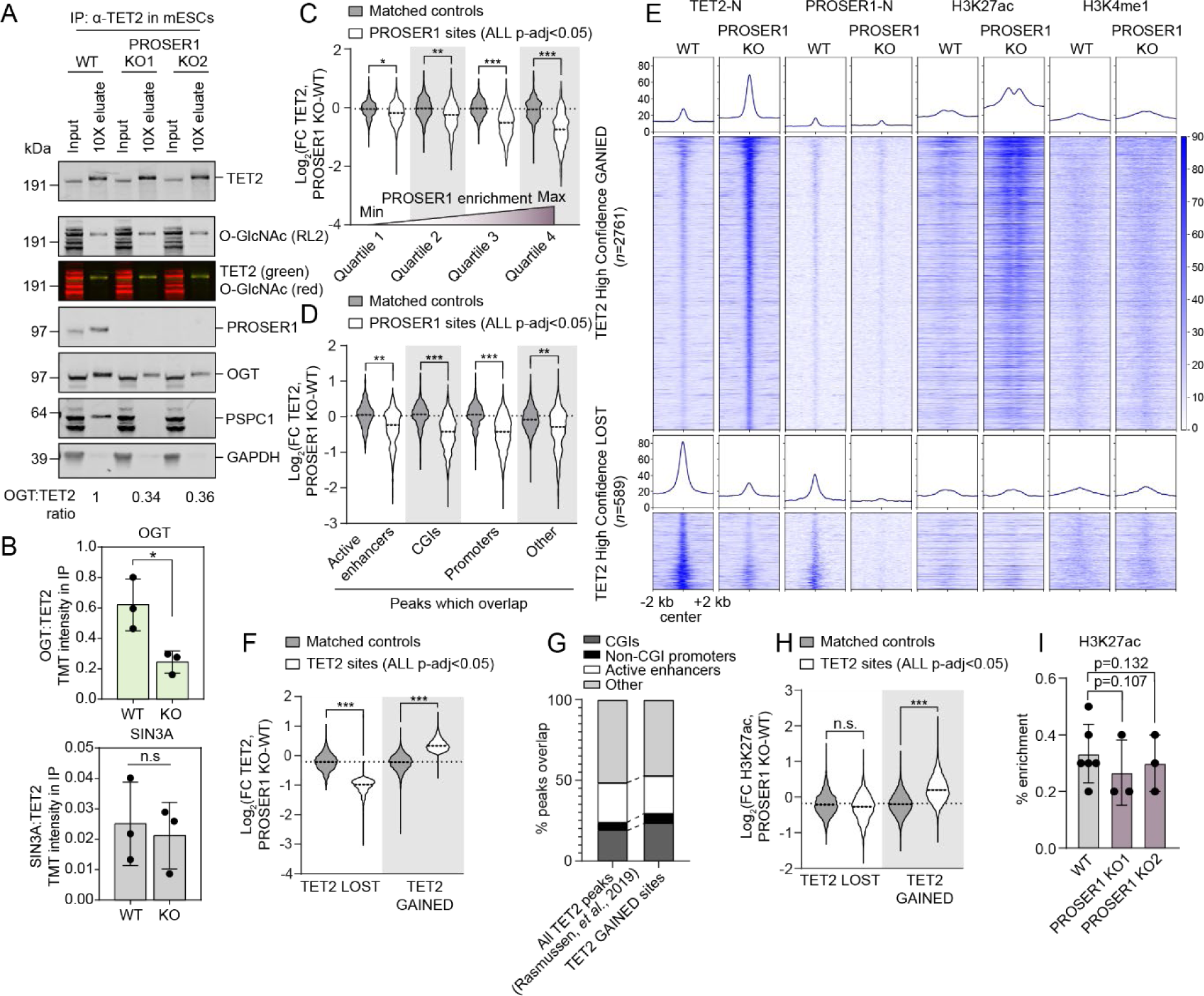
PROSER1 loss disrupts TOPD complexes and alters TET2 genome-wide chromatin binding. [A] Western blot of input and 10X concentration eluates from α-TET2 IP on lysates from WT and PROSER1 KO1 and 2 mESCs. The intensity of OGT in each eluate was normalized to the intensity of TET2 in the same eluate and the ratios in each IP provided beneath the figure. A slight difference in apparent molecular weight was observed between inputs and eluates owing to different buffer compositions. * indicates a non-specific reactive band. [B] Ratios of OGT:TET2 and SIN3A:TET2 in TMT-labelled quantitative mass spectrometry analysis of TET2 immunoprecipitate in WT and PROSER1 KO mESCs. TMT reporter intensity was quantified from 3 biological replicates. Error bars represent the mean ±S.D. * p<0.05, unpaired two-tailed *t*-test with Welch’s correction. [C] Fold change in TET2 binding based on TET2 normalized read counts within PROSER1 peaks or matched control regions upon PROSER1 loss. PROSER1 peaks were sorted by Log_2_(fold change in PROSER1, KO-WT) and divided into equal-sized quartiles (white). Controls (gray) were generated for each quartile. The effect sizes of binding loss compared to matched control regions were measured with Cohen’s d. * d>0.3 (small effect), ** d>0.6 (medium effect), *** d>0.9 (large effect). ns, non-significant. [D] As C, but TET2 normalized read counts are shown at PROSER1 sites within different genomic regions (white) as specified below the plot, or controls (gray) generated for the set of PROSER1 peaks within each genomic region. [E] Heatmaps and mean values of normalized ChIP-seq signals for TET2, PROSER1, H3K27ac and H3K4me3 centered at high-confidence (p-adj<0.05, abs(fold change≥2)) sites with gain or loss of TET2 binding. Regions are ranked on p-adj values for TET2 binding between PROSER1 KO and WT. Heatmaps were generated using DeepTools software (Ramírez et al. 2014). [F] Fold change in TET2 binding based on TET2 normalized read counts within all significant (p-adj<0.05) differential sites with gain or loss of TET2 binding (white) or matched control regions (gray) upon PROSER1 loss. The effect sizes of binding loss compared to matched control regions were measured with Cohen’s d. * d>0.3 (small effect), ** d>0.6 (medium effect), *** d>0.9 (large effect). ns, non-significant. [G] Percentage of all TET2 peaks (Rasmussen et al. 2019) or all sites with significant (p-adj<0.05) gain of TET2 binding that overlap with genomic regions as specified above the plot. [H] As F, but showing fold change in H3K27ac. [I] Global enrichment of H3K27ac in WT and PROSER1 KO1 and 2 mESCs as measured by quantitative MS. *n=*3 biological replicates of PROSER1 KO lines and 6 of WT. Error bars represent the mean ±SD. Statistical significance was measured by unpaired two-tailed *t*-test with Welch’s correction.

To investigate the association between PROSER1, TOPD recruitment to chromatin, and the activation status of PROSER1-bound regions, we then performed ChIP-seq for TET2 as well as the histone marks H3K4me1 and H3K27ac in wildtype and PROSER1 knockout mESCs (Fig. S3B). Differential enrichment analysis within high-confidence PROSER1 binding sites revealed that loss of PROSER1 correlates with reduced TET2 chromatin occupancy, while enrichment of H3K4me1 or H3K27ac does not change in response to loss of PROSER1 within the same regions (Fig. 3C, S3C, D, E). We observed that TET2 binding loss is not limited to PROSER1 binding sites within specific regulatory domains but rather is seen across active enhancers, promoters and CpG islands (CGIs) (Fig. 3D). In contrast to the reduction of TET2 binding at PROSER1-bound genomic regions, we also observed a significant number of sites with increased TET2 occupancy upon knockout of PROSER1 (Fig. 3E, F). These regions do not show evidence of PROSER1 binding in wildtype cells (Fig. 3E, and S3E), or skewing towards increased association with specific genomic regions compared to TET2 binding sites in wildtype cells (Fig. 3G). However, these regions were linked to increases in H3K27ac upon PROSER1 knockout (Fig. 3E, 3H, S3F), possibly via interactions of TET2 with the histone acetyltransferase P300, as reported previously (Zhang et al. 2017). Of note, we did not observe global differences in P300-directed histone acetylation via histone mass spectrometry, nor local differences at matched control regions which were not bound by TET2, suggesting that increased H3K27ac deposition is largely restricted to sites associated with increased TET2 binding (Fig. 3H, 3I, and S3G). Collectively, our results demonstrate that PROSER1 is required for the stability of TOPD protein complexes, and that loss of PROSER1 alters the recruitment of TET2 to chromatin.

### PROSER1 knockout unleashes TET catalytic activity and causes widespread DNA demethylation and de-silencing of endogenous retroviruses

To determine the effect of PROSER1 loss on DNA methylation, we harvested genomic DNA from two independent PROSER1 knockout mESC lines and quantified global levels of 5hmdC and 5mdC by mass spectrometry. We observed a decrease in global 5mdC levels, as well as slightly elevated levels of genomic 5hmdC - the major product of TET catalytic activity (Fig. 4A). To investigate which regions are affected by increased TET activity, we generated base-resolution DNA methylation profiles using Enzymatic Methyl-Sequencing (EM-seq) in wildtype ESCs (WT), PROSER1 knockout (PROSER1 KO), and PROSER1 knockout cells that were engineered to re-express full-length FLAG-tagged PROSER1 (KO+Rescue) (Fig. 4B). Mapping of EM-seq reads to the mouse genome allowed quantification of cytosine modification states at ∼17.5 million CpG sites with at least 10X coverage in all three genotypes. Consistent with our mass spectrometry results, we observed a decrease in DNA methylation in 10 kb windows across the entire genome in PROSER1 KO cells, while reintroduction of PROSER1 restored methylation to wildtype levels (Fig. 4C). This widespread DNA hypomethylation was also observed when comparing average DNA methylation levels in diverse genomic regions including heterochromatin, gene bodies, active enhancers and non-CGI promoters (Fig. 4D), though it was noted that regions generally depleted of DNA methylation, such as CGIs and bivalent promoters, were unchanged (Fig. S4A). We furthermore found significant DNA hypomethylation at sites associated with increased H3K27ac deposition identified previously (Fig. 4E and S3B). Importantly, expression and protein copy numbers of the major DNA methylation effectors (DNMT1/UHRF1, DNMT3A/B, TET1 and TET2) were largely unchanged (Fig. 4F, S4B). This implies that DNA methylation changes are a direct consequence of altered TET activity in PROSER1 knockout cells rather than a result of a general disruption of DNA methylation maintenance machinery.

**Figure 4.**
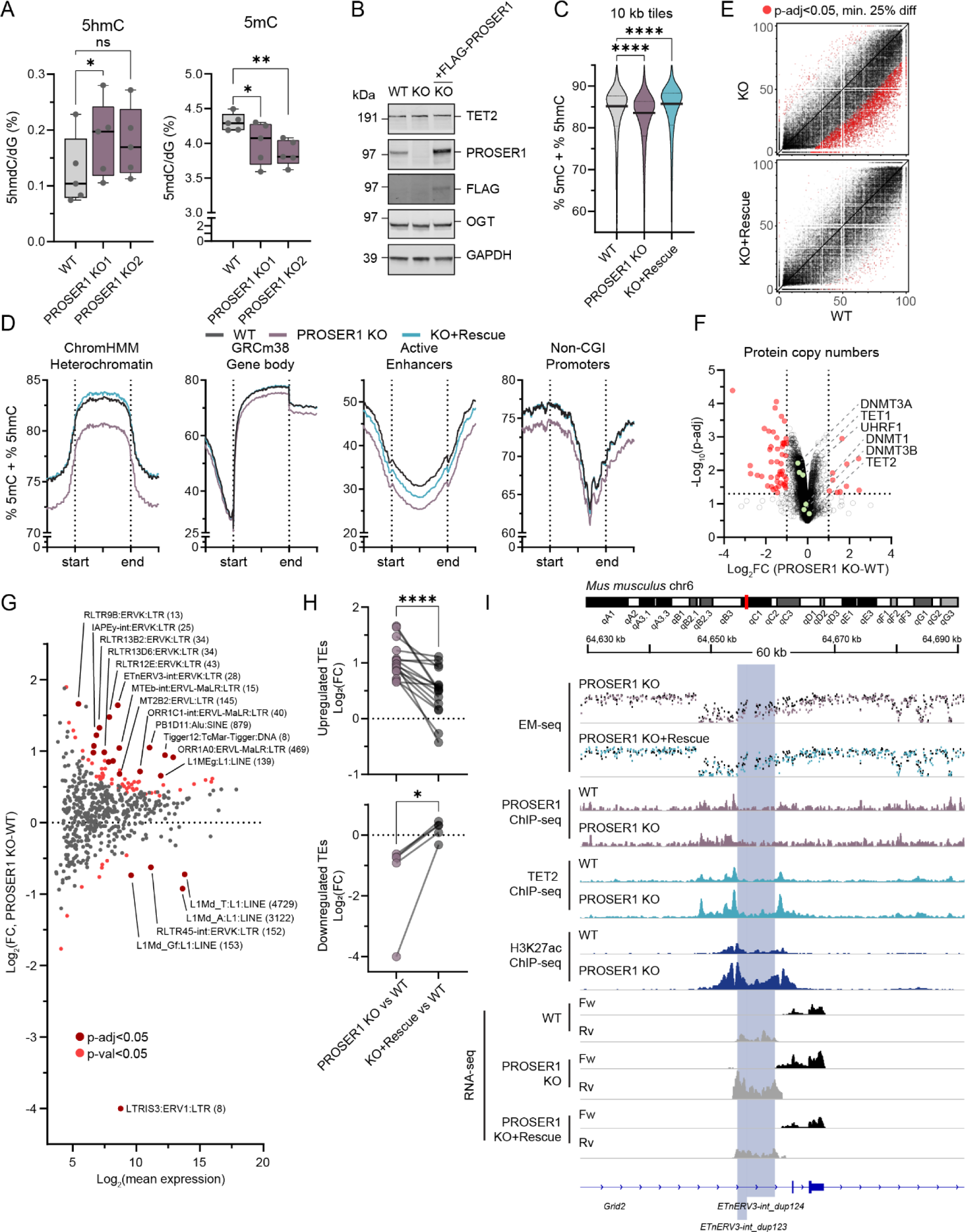
PROSER1 loss unleashes TET activity and causes widespread DNA demethylation and de-repression of endogenous retroviruses. [A] Quantification of 5hmdC and 5mdC by LC-MS/MS in genomic DNA harvested from WT, and PROSER1 KO1 and 2 mESCs. Each symbol represents a sample harvested and processed independently (*n=*5 biological replicates with each 2 technical replicates) from each cell line. Statistical significance was measured by paired two-tailed *t* test (paired by same day of harvest due to coordinated fluctuations in 5hmC). ns indicates not significant (* p<0.05, ** p<0.01). [B] Western blot of lysates prepared from WT, PROSER1 KO and PROSER1 KO+Rescue mESCs. GAPDH was probed as a loading control. [C] Violin plot showing average DNA methylation in 10 kb tiles for CpG sites covered by minimum 10 EM-seq reads in all samples in WT, PROSER1 KO and PROSER KO+Rescue mESCs. Lines on violin plot represent the median and quartiles. Statistical significance was measured by Brown-Forsythe and Welch ANOVA test (**** p<0.0001). [D] Quantitation trend plots of DNA methylation quantified by EM-seq in heterochromatin, gene bodies, active enhancers and non-CGI promoters for CpG sites covered by minimum 10 EM-seq reads in all samples in WT, PROSER1 KO and PROSER1 KO+Rescue mESCs. [E] XY-scatter plot showing modification state of individual CpG sites covered by at least 10 EM-seq reads that overlap 3196 regions of increased H3K27ac deposition identified previously (Fig. S3B) in PROSER1 KO vs WT cells. Significantly differentially methylated CpG sites (p-adj<0.05, and minimum 25% difference) are highlighted in red for PROSER1 KO vs WT (above) and KO+Rescue vs WT (below). [F] Volcano plot showing protein copy numbers determined by whole-proteome MS on PROSER1 KO mESCs (*n=*3 biological replicates). Protein enrichment is compared to parallel MS in WT mESCs. Dotted lines indicate 2-fold change and p-adj 0.05. Components of the DNA methylation machinery are highlighted and identified. [G] MA-plot showing *TEtranscripts* differential expression analysis of TEs in WT and PROSER1 KO mESCs. Each dot represents a sub-family of TE elements and the number of individual elements included in the analysis (covered by at least one unique read) is shown in parentheses. Red and dark red dots indicate significantly differentially expressed TE families at thresholds of p-value<0.05 and p-adj<0.05, respectively. [H] Symbol and line plots comparing expression of TE families in PROSER1 KO vs WT and PROSER1 KO+Rescue vs WT. Plots include TE families found to be differentially expressed (p-adj<0.05) between WT and PROSER1 KO in [G] and split to show upregulated (above) and downregulated (below) TE families. Statistical significance was measured by paired two-tailed *t* test (* p<0.05, **** p<0.0001). [I] Tracks showing the region surrounding *ETnERV3-int_dup123* and *124* (highlighted in blue) which gain TET2 binding and H3K27ac and become DNA hypomethylated and de-silenced in the absence of PROSER1. Data in the top two panels represent pooled EM-seq methylation calls. Data from WT mESCs are shown in black and data from PROSER1 KO and PROSER1 KO+Rescue are overlaid in purple or blue, respectively. Below, tracks represent ChIP-seq coverage in PROSER1 ChIP-seq (*n=*2 biological replicates), and TET2 or H3K27ac ChIP-seq (*n=*3 biological replicates). The last 6 tracks represent coverage of forward (Fw) or reverse (Rv) transcripts identified by RNA-seq (*n=*2 biological replicates).

Dual pharmacological inhibition of DNA methylation enzymes and histone deacetylases causes DNA hypomethylation and increased histone acetylation - reminiscent of changes observed upon PROSER1 knockout – and synergizes to cause de-silencing of transposable elements (TEs) (Brocks et al. 2017; Daskalakis et al. 2018; Goyal et al. 2023; Cusack et al. 2020). We therefore used a combination of RNA-seq and *TEtranscripts* - an analysis pipeline designed to handle reads that map to multiple locations in the genome - to assign multi-mapping reads to specific TE families and analyze their activity. Loss of PROSER1 led to an increase in transcription of multiple families of long terminal repeat (LTR)-containing endogenous retroviral (ERV) elements, such as ERVK, ERVL, and ERVL-MaLR, whereas expression of the non-LTR L1Md retrotransposons was mildly reduced (Fig. 4G). To understand if this deregulation correlated with loss of PROSER1, we assessed transcript levels of the differentially expressed TE families upon re-expression of PROSER1. Consistent with the observed restoration of DNA methylation levels described above, expression of differentially expressed TE families (Fig. 4H) – as well as expression of individual TE elements identified solely based on uniquely mapped reads (Fig. 4I and S4C) – were restored to near-wildtype levels upon reintroduction of full-length PROSER1. Collectively, our findings demonstrate that PROSER1 safeguards against genome-wide DNA demethylation and aberrant activation of endogenous retroviruses.

### Defective recruitment of TET2 to developmental genes upon PROSER1 loss leads to their subsequent dysregulation during neuronal differentiation

To determine the direct effect of altered TET2 chromatin binding, we quantified DNA methylation changes in the high-confidence TET2 differentially bound sites (p-adj<0.05, abs(fold change)≥2) identified in PROSER1 knockout cells (Fig. 3E). In contrast to regions with gained TET2 binding - which mirrored the genome wide DNA hypomethylation - sites with reduced TET2 binding were instead correlated with increased levels of DNA methylation (Fig. 5A). Further analysis identified that nearly a third of sites with reduced TET2 binding exhibited a significant rise in average DNA methylation (p-adj<0.05, minimum 3 CpG per site) (Fig. 5B). To investigate potential effects on gene expression, we identified enriched gene ontology terms in the subset of 866 genes whose regulatory domains (defined in GREAT as “Basal plus extension” (McLean et al. 2010)) overlapped with sites with reduced TET2 binding. This analysis revealed significant associations with early development processes, such as nervous and skeletal system development, and was linked to mouse knockout phenotypes exhibiting craniofacial abnormalities including eye defects (Fig. 5C). In contrast, genes whose regulatory domains overlapped with sites with increased TET2 binding were largely linked to phenotypes associated with abnormal hematopoietic differentiation (Fig. S5A).

**Figure 5.**
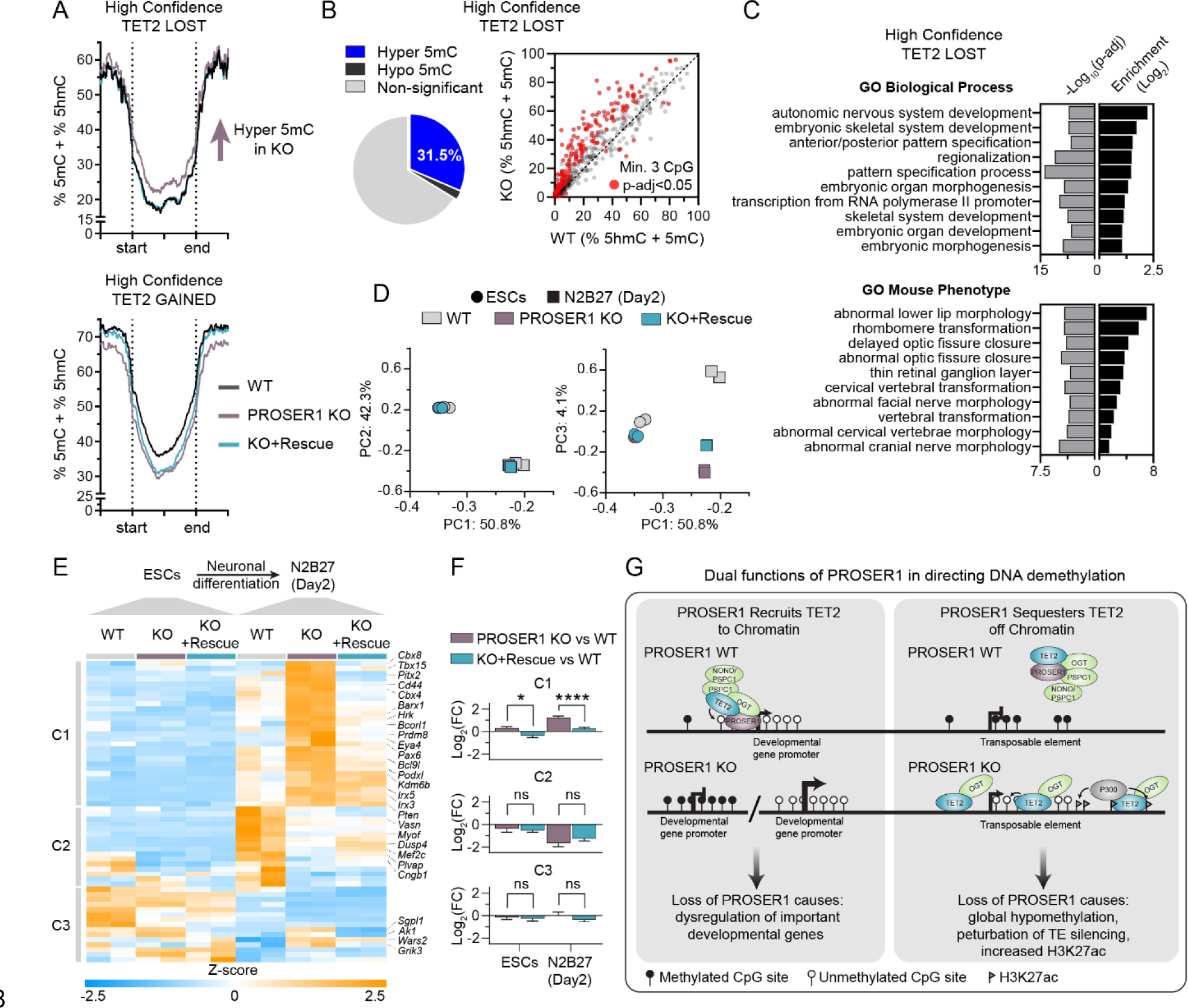
Defective recruitment of TET2 to developmental genes upon PROSER1 loss leads to their subsequent dysregulation during neuronal differentiation. [A] Quantitation trend plots of DNA methylation for CpG sites covered by minimum 10 EM-seq reads in all samples in WT, PROSER1 KO and PROSER1 KO+Rescue mESCs at high-confidence (p-adj<0.05, Abs(fold change≥2)) sites with loss (above) or gain (below) of TET2 binding upon PROSER1 KO. [B] Pie chart (left) and XY-scatter (right) showing average DNA methylation within high-confidence sites with loss of TET2 binding upon PROSER1 KO. Significantly (p-adj<0.05) hypermethylated or hypomethylated sites (min. 3 CpG per site, min. 10 reads per CpG) are indicated. [C] Bar charts showing enriched gene ontology (GO) terms identified by GREAT (McLean et al. 2010) for genes with a regulatory domain overlapping high-confidence sites with loss of TET2 binding upon PROSER1 KO. [D] PCA plot of RNA-seq data. PCA was carried out using Z-scores of the top 2500 variable genes in WT, PROSER1 KO1 and PROSER1 KO+Rescue cells in the mESC state (circular markers) or after 2 days of differentiation towards neuronal progenitors (square markers). [E] Heatmap showing Z-scores of genes differentially expressed at day 2 of neuronal differentiation between PROSER1 KO and WT (p-adj<0.05) with regulatory domains overlapping high-confidence sites with loss of TET2 binding upon PROSER1 KO in mESCs. Hierarchical clustering identified three distinct clusters C1-C3 as indicated. Gene symbols for selected rows in the heatmap are given on the right of the plot. See sup. table S7 for full list of differentially expressed genes associated with loss of TET2 binding. [F] Bar charts showing average Log_2_ fold change for each individual cluster defined in [E] for PROSER1 KO vs WT and KO+Rescue vs WT in mESCs and after 2 days of neuronal differentiation respectively. Statistical significance was measured by Brown-Forsythe and Welch ANOVA test (* p<0.05, **** p<0.0001). [G] Schematic illustrating the dual roles of PROSER1 in directing TET function and DNA demethylation.

To directly assess the effect of PROSER1 knockout during mESC differentiation, we analysed gene expression in self-renewing mESC lines and in cells differentiated for 2 days towards neuronal lineages (N2B27 monolayer differentiation system). Initial inspection of RNA-seq results revealed that all cell lines successfully exited pluripotency and upregulated markers of early neuronal differentiation (Fig. S5B). Consistent with this, principal component analysis (PCA) revealed differentiation state as the main factor driving variation in the samples (PC1: 50.8% and PC2: 42.3%). In addition, we identified a minor component (PC3: 4.1%) that correlated with PROSER1 expression, and clearly separated samples cultured for 2 days in N2B27 medium (Fig. 5D). We therefore examined if the reduced TET2 chromatin binding we observed in PROSER1 knockout cells could be linked to gene expression changes at this developmental stage. To do this, we focused on a subset of 60 genes that i) had a regulatory domain overlapping sites of reduced TET2 binding in mESCs, and ii) were differentially expressed (p-adj<0.05) between wildtype and PROSER1 knockout cells upon 2 days of neuronal differentiation (Fig. 5E and sup. table S7). Contingency analysis revealed significant enrichment of this subset of genes (both increased and decreased in knockout) compared to the subset of differentially expressed genes not associated with reduced TET2 binding sites (7.43% vs 3.91% of total, p-value<0.0001, Fisher’s exact test). Hierarchical clustering identified three clusters (C1-C3) with different expression patterns broadly classified as up-, down- or mixed regulation in PROSER1 knockout cells, respectively (Fig. 5E). The observed increases and decreases in transcript levels upon PROSER1 depletion suggest that TOPD complexes possess both activating and repressive regulatory capacities, the specific outcome of which is likely influenced by genomic location and developmental stage. Importantly, comparison of Log_2_ fold changes in PROSER1 KO vs WT, and KO+Rescue vs WT showed that re-introduction of PROSER1 restored expression of many of these genes – particularly in C1 – to near-wildtype levels (Fig. 5F). Genes co-bound by PROSER1/TET2 and whose expression correlated with PROSER1 expression included important developmental regulators such as homeobox genes (e.g. *Pitx2, Barx1,* and *Pax6*), epigenetic regulators (e.g. *Cbx4, Cbx8,* and *Prdm8*), and transcription factors (e.g. *Tbx15* and *Mef2c*) implying that PROSER1 plays a regulatory role in their function during early embryonic development (Fig. S5C, D). We furthermore noted significant deregulation of Eyes absent homolog 4 (*Eya4*) – a PROSER1/TET2 co-bound target gene – that is crucial for eye, heart and sensorineural development (Fig. S5C, D) (Tadjuidje and Hegde 2013). Collectively, deregulation of these genes may underlie some, if not all, of the neurodevelopmental defects observed upon loss of PROSER1 in mice and humans.

In summary, our results are consistent with a dual role for PROSER1 in directing DNA demethylation in early development (Fig. 5G). We show that PROSER1 is a pan-TET interactor that promotes the assembly of TOPD protein complexes and facilitates their recruitment to chromatin in the proximity of important developmental genes. The recruitment of TET proteins – particularly TET2 – at PROSER1-bound sites maintains a lowly methylated state at regulatory regions and appropriate expression of adjacent genes during differentiation. Our findings also indicate that TOPD complexes sequester TET proteins away from other regions of the genome. When PROSER1 is depleted, TET proteins bind to additional sites, resulting in widespread DNA demethylation, and therefore global DNA hypomethylation. Interestingly, the regions with increased TET2 binding also displayed increased levels of H3K27 acetylation. This suggests that P300, a histone acetyltransferase previously shown to interact with TET2 (Zhang et al. 2017), might be recruited alongside TET2 to these same sites. The combined effects of reduced DNA methylation and increased chromatin openness (caused by P300 activity) cooperate to perturb the silencing of endogenous retroviral elements, potentially disrupting the expression of adjacent genes during differentiation.

The exact mechanism by which TOPD complexes control TET activity across the genome remains unclear. One hypothesis is that certain PROSER1 binding sites act as “sinks,” attracting TOPD complexes and TET proteins to specific genomic sites and preventing widespread, uncontrolled DNA demethylation by TET enzymes. Another interesting possibility involves the RNA-binding TOPD component PSPC1, which we have found to interact with TET2 in a PROSER1-dependent manner (Fig. 3A and S3A). DBHS proteins including PSPC1 can form higher-order oligomers (Fox et al. 2018; Knott et al. 2016, 2022), potentially functioning as RNA- or DNA-tethered condensates that sequester TOPD complexes, further regulating TET activity. Consistent with this, biochemical fractionation experiments in mESCs revealed that the ability of TET2 to fractionate with chromatin is largely dependent on both PSPC1 and its RNA binding capacity (Guallar et al. 2018). Additional work will be needed to address these possibilities in further detail.

Our findings align with prior research. Recent work revealed that introduction of a missense mutation into *Tet1* in mESCs leads to a partial disruption of TET1-OGT interaction. This results in global DNA hypomethylation, consistent with the mutated TET1 protein detaching from TOPD complexes to trigger widespread DNA demethylation (Hrit et al. 2018). Similarly, a recent study observed global DNA demethylation and TE de-repression after acute deletion of OGT in mESCs (Sepulveda et al. 2024). This strongly suggests that OGT loss disrupts TOPD complex stability, leading to the release of TET proteins and subsequent genome-wide DNA demethylation. Interestingly, the absence of TET enzymes themselves has been linked to widespread DNA demethylation, particularly in heterochromatin (López-Moyado et al. 2019). This phenomenon was proposed to stem from disruption of a shared protein complex important for both TET and DNMT enzyme recruitment. When TET enzymes are removed, this complex is disrupted, leading to the redistribution of DNMT enzymes across the genome (López-Moyado et al. 2019). Thus, in addition to the role of PROSER1 in restraining TET activity, it is interesting to speculate whether TOPD complexes may have additional roles – directly or indirectly - in regulating DNMT activity.

Pair-wise TET interactions with specific partner proteins like OGT, PSPC1, and NONO have previously been identified (Chen et al. 2013; Vella et al. 2013; Deplus et al. 2013; Guallar et al. 2018; Huang et al. 2022; Li et al. 2020). However, our results imply that these interactions do not occur in isolation but can combine to form larger multimeric TOPD complexes that have functional roles in chromatin. Interestingly, mutations in several components of TOPD complexes beyond PROSER1 have been linked to neurodevelopmental disorders. These include X-linked variants of OGT (causing Congenital Disorder of Glycosylation (OGT-CDG))(Pravata et al. 2020; Authier et al. 2024) and NONO (causing NONO-associated syndromic disorder)(Mircsof et al. 2015; Reinstein et al. 2016; Roessler et al. 2023) as well as biallelic loss of TET3 (causing Beck-Fahrner syndrome)(Beck et al. 2020; Seyama et al. 2022), all of which display syndromic developmental delay, intellectual disability and craniofacial dysmorphisms similar to features observed upon inactivation of PROSER1. While mutations in TOPD components are likely to have additional pleiotropic effects, some common features across these disorders may stem from shared disruption of TOPD complexes during development.

Chromatinopathies represent an expanding category of congenital developmental disorders arising from disruptions in chromatin function and dysregulation of the epigenome (Nava and Arboleda 2024). Mutations within genes encoding critical epigenome regulators, encompassing both core components and accessory proteins such as PROSER1, contribute to this growing list of pathologies. In this study, we show that PROSER1 plays a central role in the assembly of multi-protein chromatin-associated TET complexes that shape the DNA methylome and support gene expression. Furthermore, mice lacking PROSER1 mirror the developmental defects seen in humans with homozygous PROSER1 loss-of-function mutations. We therefore propose that developmental syndromes caused by PROSER1 mutations should be designated as a novel form of chromatinopathy and that future investigations into TOPD-related developmental syndromes should leverage the growing understanding of TET enzyme function in development and disease. Our development of a PROSER1 knockout mouse model serves both as a system to gain insight into the underlying mechanisms, and a preclinical model to explore the potential for therapeutic intervention in PROSER1-related developmental syndromes.

## MATERIALS AND METHODS

See supplementary methods for further information.

## DATA AVAILIBILITY

Raw and processed data sets are available for download at the Gene Expression Omnibus (GEO) database under the accession number GSE273517.

## COMPETING INTEREST STATEMENT

The authors declare no competing interests

## ACKNOWLEDGEMENTS

The authors thank members of the Rasmussen lab, G. Saredi, and T. Owen-Hughes for advice and discussion. We thank A. Rennie and R. Clarke in the Flow Cytometry unit, C. Gillian in the Biological Resource Unit, A. Atrih and C. Rogers in Fingerprints proteomics facility at University of Dundee as well as A. Tavares and C. Corral in the imaging facility at the University of Edinburgh for technical assistance. Work in the Rasmussen lab was funded by a Cancer Research UK fellowship (C66224/A27092) and through support by University of Dundee. X.L. was funded through a personal scholarship provided by the China Scholarship Council (CSC). Work in the Hajkova laboratory is supported by MRC funding 815 (MC_US_A652_5PY70) and an ERC grant (ERC-CoG-648879–dynamicmodifications). The Sidoli lab gratefully acknowledges, for funding, the Hevolution Foundation (AFAR), the Einstein-Mount Sinai Diabetes center, and the NIH Office of the Director (S10OD030286). The authors thank the dedicated team behind the European Galaxy server (UseGalaxy.eu), supported by the German Federal Ministry of Education and Research grant 031L0101C and de.NBI-epi.

## AUTHOR CONTRIBUTIONS

A.F and K.D.R. conceived the study. A.F., E.V.K., X.L., and K.D.R. carried out experiments. A.Z., and D.O.C. generated the PROSER1 knockout mouse model. Z.H. and P.H. conducted 5hmdC/5mdC mass spectrometry experiments and analysis. S.S. and S.S. conducted histone mod mass spectrometry experiments and analysis. A.J.B. analyzed mass spectrometry data. A.F. and K.D.R. performed the bioinformatics analysis. A.F. and K.D.R. wrote the original draft of the manuscript followed by review and editing by all authors. K.D.R. supervized the study and acquired funding.

---

**Figure S1.**
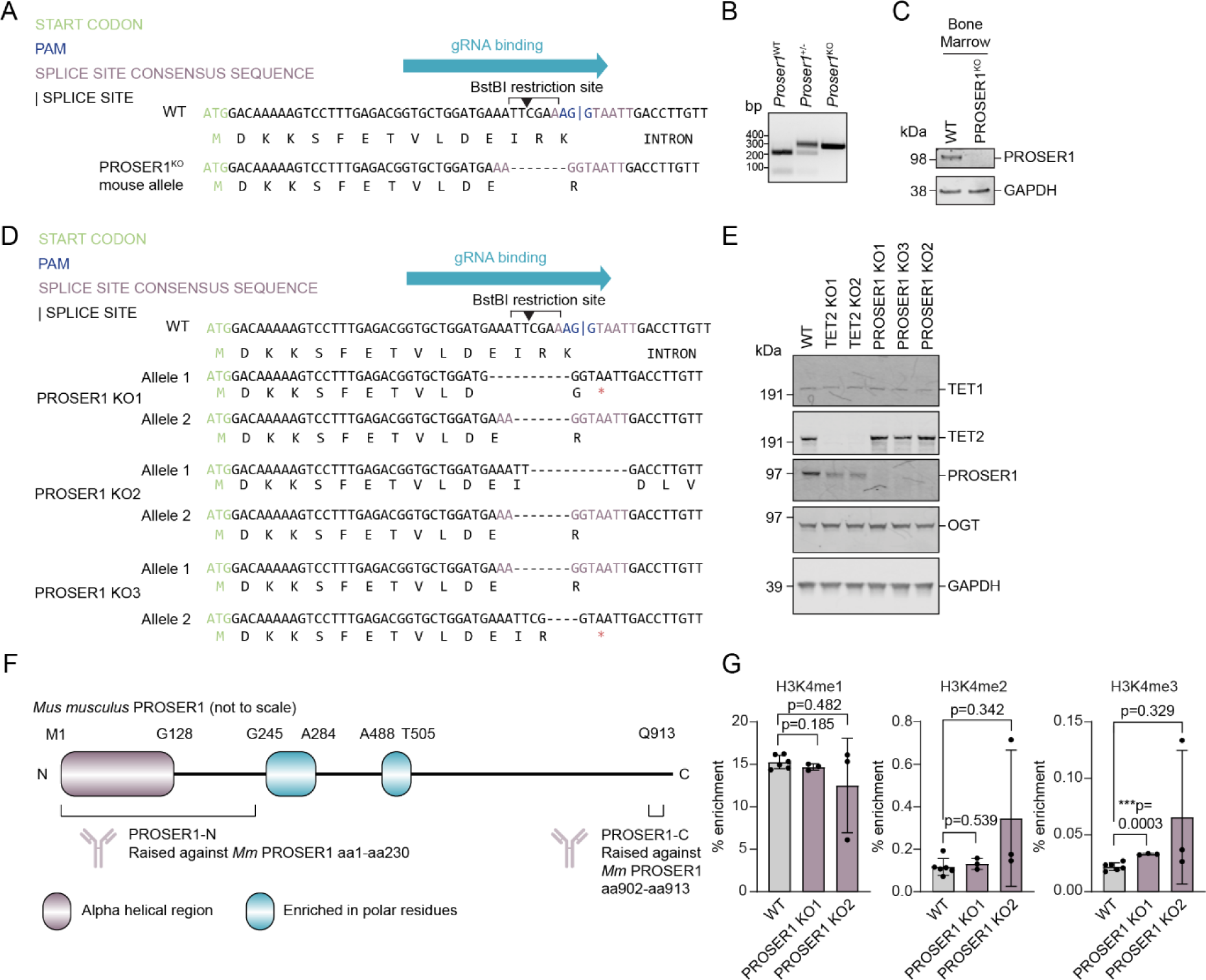
[A] DNA and amino acid sequences of the region surrounding the PROSER1 gRNA binding site in WT and PROSER1^KO^ mice. Green text indicates the start codon, blue the PAM sequence, purple the splice-site consensus sequence. The BstBI restriction site is indicated above the DNA sequence. PROSER1^KO^ mice contains a 7bp deletion that disrupts the reading frame and introduces a premature stop codon. [B] Representative agarose gel showing genotyping of a wildtype (WT), PROSER1 heterozygous (+/-), and knockout (KO) mouse. bp indicates base pairs [C] Western blot of lysates from the bone marrow of a WT or PROSER1^KO^ mouse. GAPDH was probed as a loading control. [D] As in A, but in three clonal PROSER1 KO mESC lines. When an allele has no purple regions this indicates that the splice-site consensus sequence is lost. Premature stop codons are indicated with red asterisks. Both alleles in these three clones either had a frameshift mutation or lost the splice-site consensus sequence. [B] Western blot of lysates from WT, TET2 KO or PROSER1 KO clonal mESCs. GAPDH was probed as a loading control. [C] Illustration of *Mus musculus* PROSER1 (not to scale), with the binding regions of our PROSER1-N and PROSER1-C antibodies indicated below. The single letter amino acid codes and the residue numbers at which the indicated regions begin and end are given above. [D] Global enrichment of H3K4me1, 2 or 3 in WT and PROSER1 KO1 and 2 mESCs as measured by quantitative MS. Error bars represent the mean ±SD. Statistical significance was measured by unpaired two-tailed *t*-test with Welch’s correction.

**Figure S2.**
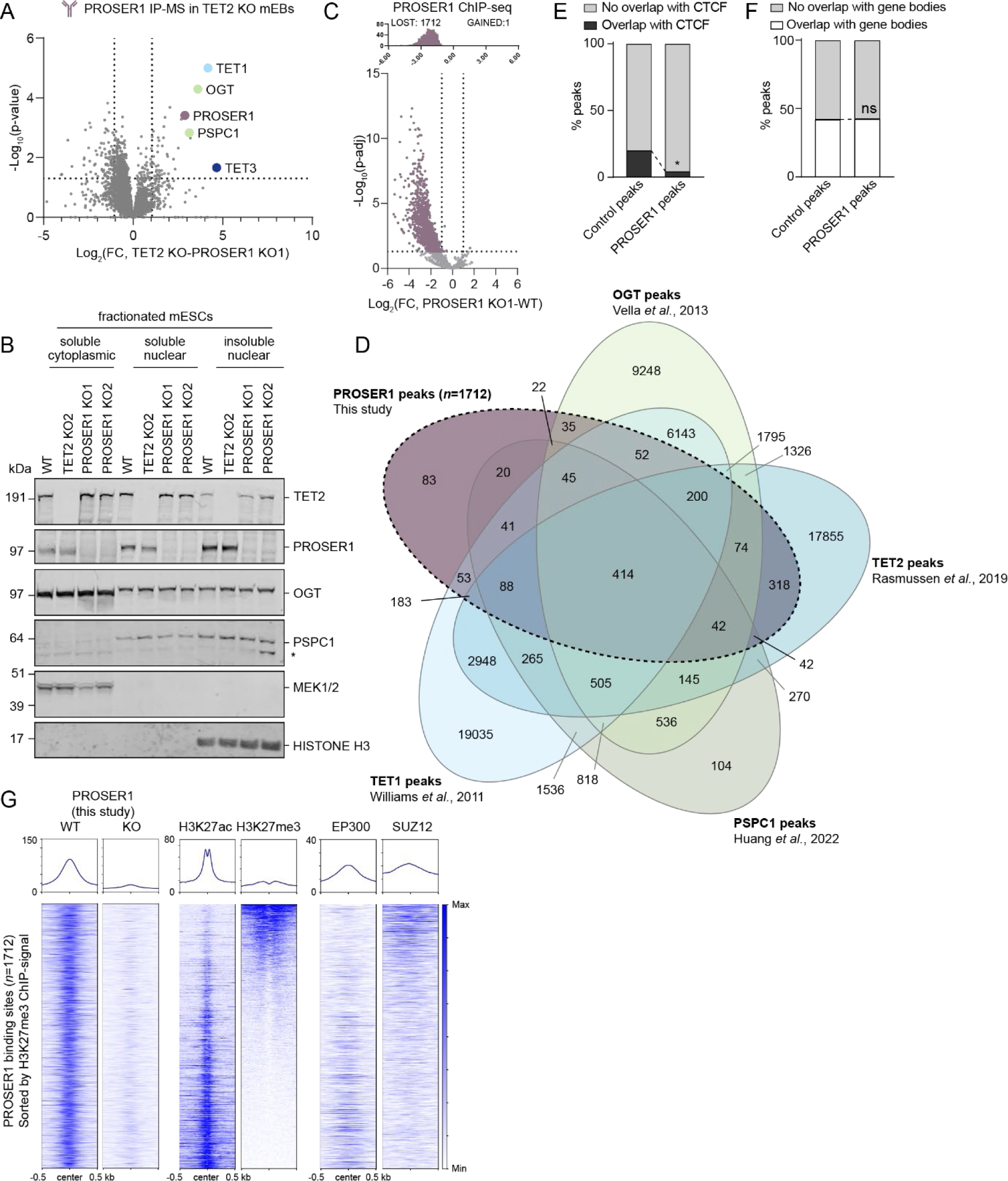
[A] Volcano plot showing protein hits from α-PROSER1-C immunoprecipitation and mass spectrometry (IP-MS) in TET2 KO mEBs (*n=*3 biological replicates). Protein enrichment is compared to parallel IP-MS in PROSER1 KO mEBs. Dotted lines indicate 2-fold change and p-value 0.05. Proteins of interest are highlighted and identified. See sup. table S5 for full list of enriched proteins. [B] Western blot of fractionated WT, TET2 KO and PROSER1 KO1 and 2 mESCs. MEK1/2 and histone H3 were run as fractionation controls for the cytoplasmic and chromatin fractions, respectively. * indicates a non-specific reactive band. [C] Volcano plot and frequency histogram showing changes in PROSER1-chromatin binding identified by PROSER1-N ChIP-seq in WT and PROSER1 KO1 mESCs (*n=*2 biological replicates). Dotted lines indicate 2-fold change and p-adj 0.05. PROSER1-chromatin binding sites where p-adj<0.05 and abs(fold change≥2) are highlighted in purple and their number given. [D] Venn diagram showing overlap between PROSER1, OGT, TET2, PSPC1 and TET1 peaks in mESCs. The number of overlapping peaks in each combination of peaksets is indicated. All peaksets but PROSER1-N peaks were generated from publicly available data (Huang et al. 2022; Rasmussen et al. 2019; Vella et al. 2013; Williams et al. 2011). [E] Percentage of PROSER1-N peaks or matched controls that overlap with CTCF binding. CTCF binding sites were obtained from publicly available data (Song et al. 2022). * p<0.0001, two-tailed Fisher’s exact test. [F] As E, but overlapping gene bodies. [G] Heatmaps and mean values of normalized ChIP-seq signals for PROSER1-N and activating (P300 and H3K27ac) and repressive (H3K27me3 and SUZ12) chromatin features centered at 1712 high-confidence PROSER1 binding sites in mESCs. P300 and SUZ12 enrichment were generated from (Wang et al. 2017) and (Højfeldt et al. 2018) respectively, and regions were ranked based on the H3K27me3 ChIP-seq signal. Heatmaps were generated using DeepTools software (Ramírez et al. 2014).

**Figure S3.**
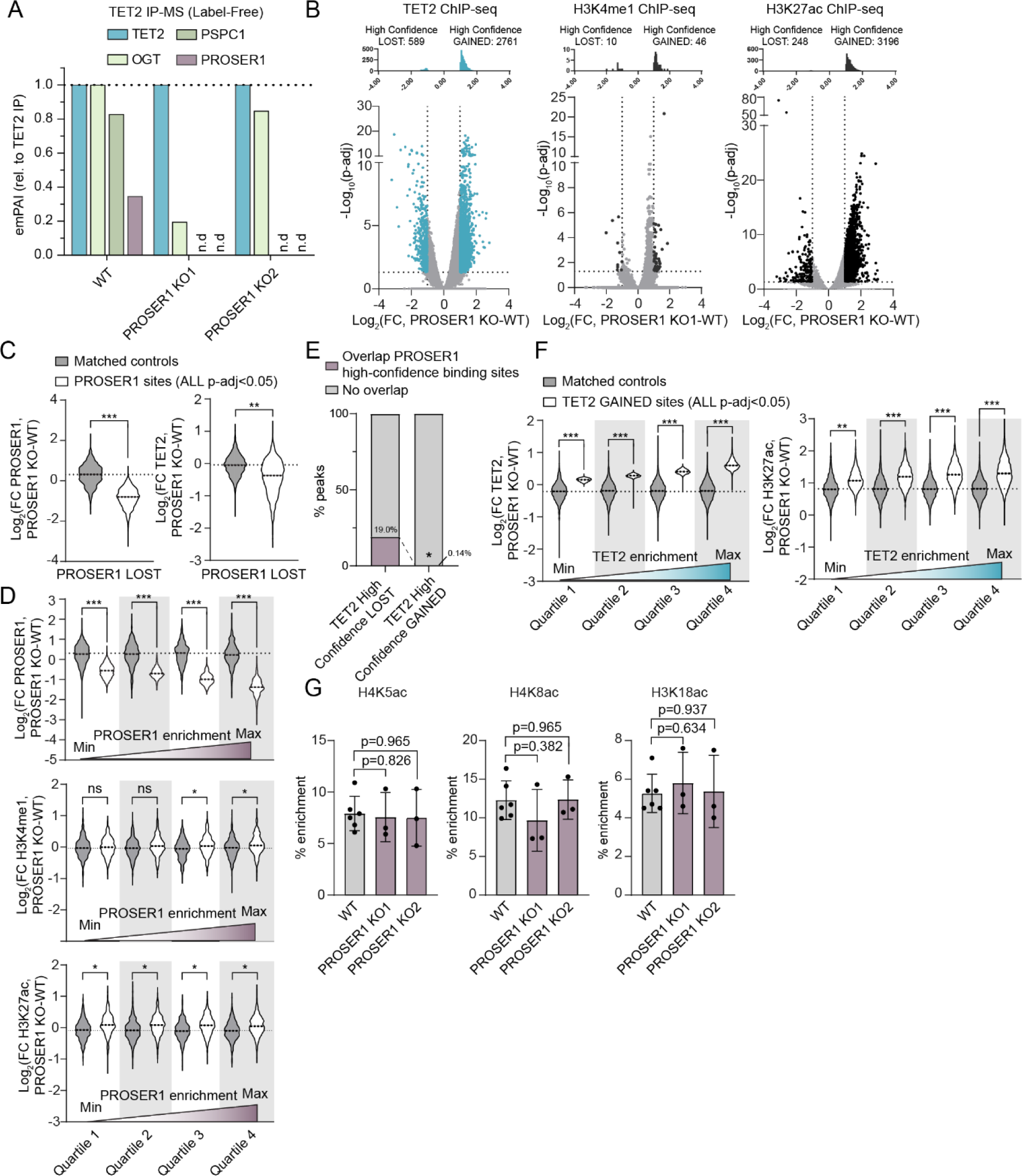
[A] Estimated protein abundance indicated by exponentially modified protein abundance index (emPAI) of TET2, OGT, PSPC1 and PROSER1 in TET2 IP-MS normalized to TET2 IP efficiencies in each experiment (*n=*1 biological replicate). n.d., not detected. [B] Volcano plot and frequency histograms showing changes in TET2 (left), H3K4me1 (center) or H3K27ac (right) enrichment on chromatin identified by ChIP-seq in WT and PROSER1 KO1 mESCs (*n=*3 biological replicates). Dotted lines indicate 2-fold change and p-adj 0.05. Sites where p-adj<0.05 and Abs(fold change≥2) are highlighted in blue or black (for TET2 or H3K4me1 and H3K27ac, respectively) and their number is given. [C] Fold change in PROSER1 binding (left) or TET2 binding (right) based on normalized read counts within PROSER1 DOWN and PROSER1 UP sites (p-adj<0.05) (white) or matched control regions (gray) upon PROSER1 loss. The effect sizes of binding loss compared to matched control regions were measured with Cohen’s d. * d>0.3 (small effect), ** d>0.6 (medium effect), *** d>0.9 (large effect). ns, not-significant. [D] Fold change in PROSER1 binding (above) or enrichment of H3K4me1 (center) or H3K27ac (below) within PROSER1 peaks or matched control regions upon PROSER1 loss. PROSER1 peaks were sorted by Log_2_(fold change in PROSER1, KO-WT) and divided into equal-sized quartiles (white). Controls (gray) were generated for each quartile. The effect sizes of binding loss compared to matched control regions were measured with Cohen’s d. * d>0.3 (small effect), ** d>0.6 (medium effect), *** d>0.9 (large effect). ns, non-significant. [E] Percentage of PROSER1 peaks that overlap with high-confidence (p-adj<0.05, Abs(fold change≥2)) sites with gain or loss of TET2 binding. * p<0.0001, two-tailed Fisher’s exact test. [F] As D, but fold change in TET2 binding (left) or H3K27ac enrichment (right) at quartiles of all sites with significant (p-adj<0.05) gain of TET2 binding. [G] Global enrichment of H3K5ac (left), H4K8ac (center) and H3K18ac (right) in WT and PROSER1 KO1 and 2 mESCs as measured by quantitative MS. *n=*3 biological replicates of PROSER1 KO lines and 6 of WT. Error bars represent the mean ±SD. Statistical significance was measured by unpaired two-tailed *t*-test with Welch’s correction.

**Figure S4.**
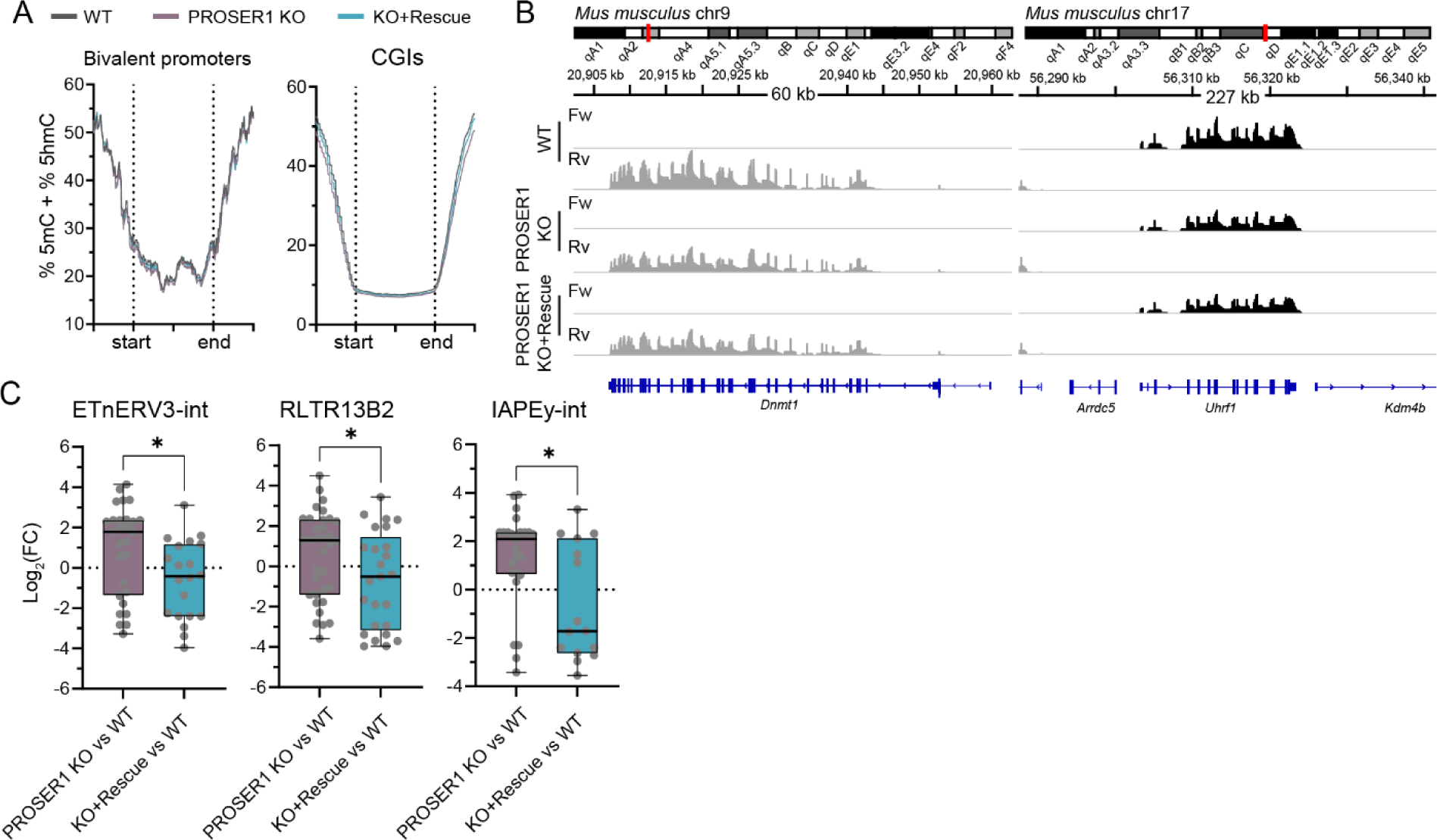
[A] Quantitation trend plots (window size 50 bp, step 50 bp, 1 kb flanking sequence) of DNA methylation quantified by EM-seq in bivalent promoters and CGIs for CpG sites covered by minimum 10 EM-seq reads in all samples in WT, PROSER1 KO and PROSER1 KO+Rescue mESCs. [B] Coverage of forward (Fw) or reverse (Rv) transcripts from RNA-seq (*n=*2 biological replicates) in WT, PROSER1 KO and KO+Rescue mESCs at the regions surrounding *Dnmt1* (left) and *Uhrf1* (right). [C] Box plots showing *TEtranscripts* differential expression analysis of TEs. Dots represent expression of individual TEs whose expression was quantified using only uniquely mapped RNA-seq reads. The box plots compare differential expression of TEs in PROSER1 KO vs WT and PROSER1 KO+Rescue vs WT for ETnERV3-int (left), RLTR13B2 (middle), and IAPEy-int (right) families of endogenous retroviral elements, all found to be significantly deregulated upon PROSER1 KO. Statistical significance was measured by unpaired two-tailed *t*-test with Welch’s correction (* p<0.05).

**Figure S5.**
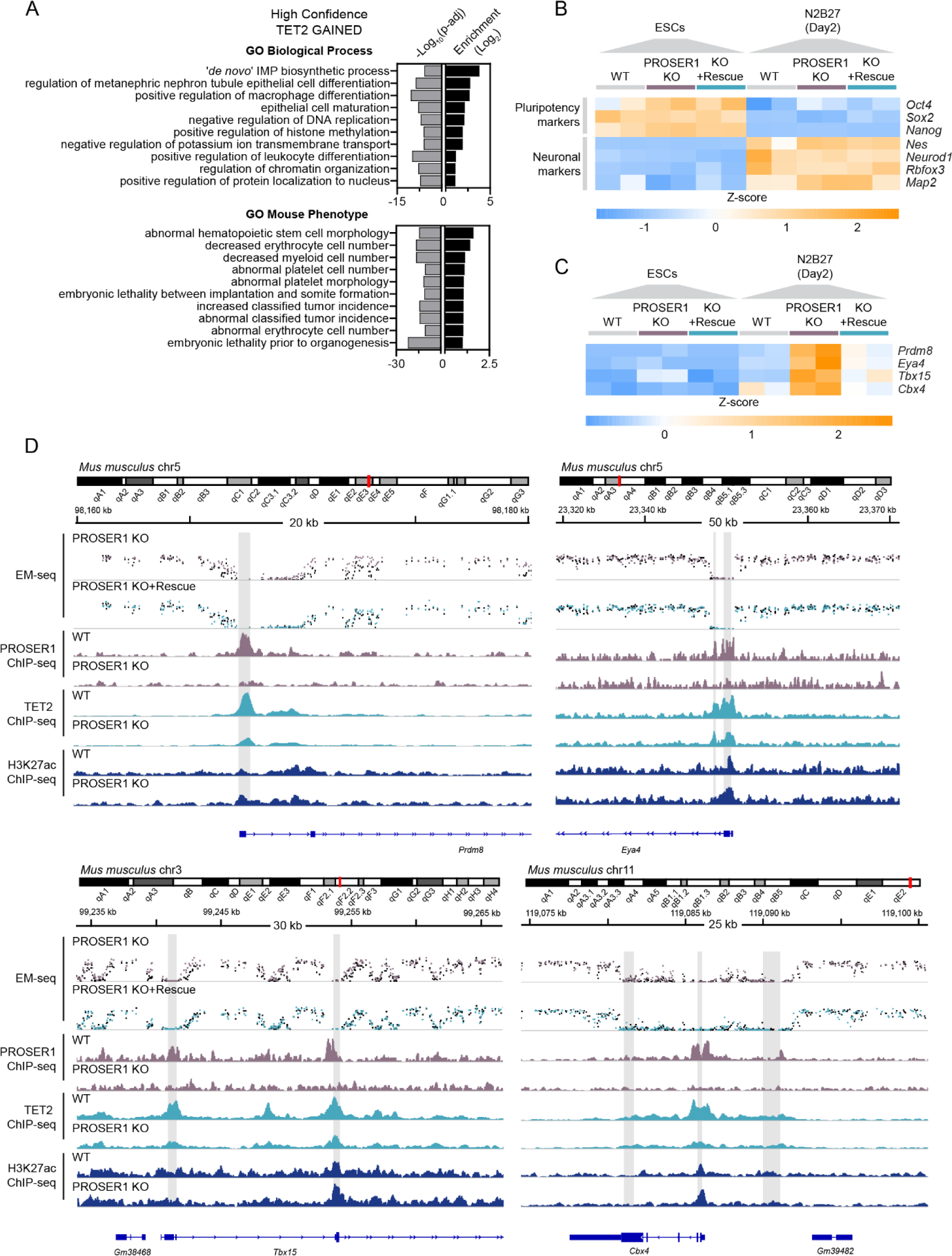
[A] Bar charts showing enriched gene ontology (GO) terms identified by GREAT (McLean et al. 2010) for genes with regulatory domain overlapping high-confidence sites with gain of TET2 binding upon PROSER1 KO. [B] Heatmap showing the changes in expression (Z-score of normalized read counts from RNA-seq) of key pluripotency markers (above), and neuronal markers (below) in WT, PROSER1 KO and PROSER1 KO+Rescue mESCs (left) or cells subjected to monolayer differentiation in N2B27 medium (right). Gene symbols for each row in the heatmap are given on the right of the plot. [C] As in B, but for selected developmental genes co-bound by PROSER1 and TET2 whose expression correlated with PROSER1 expression. [D] Tracks showing the region surrounding the developmental genes *Prdm8*, *Eya4*, *Tbx15*, and *Cbx4*. Data in the top two panels represent pooled EM-seq methylation calls. Data from WT mESCs are shown in black and data from PROSER1 KO and PROSER1 KO+Rescue are overlaid in purple or blue, respectively. Below, tracks represent ChIP-seq coverage in PROSER1 ChIP-seq (*n=*2 biological replicates), and TET2 or H3K27ac ChIP-seq (*n=*3 biological replicates). CGIs are indicated in gray.

